# End-to-end neural system identification with neural information flow

**DOI:** 10.1101/553255

**Authors:** K. Seeliger, L. Ambrogioni, Y. Güçlütürk, U. Güçlü, M. A. J. van Gerven

**Author notes:** Corresponding author (MvG). These authors contributed equally to this work (K. Seeliger, L. Ambrogioni).

## Abstract

Neural information flow (NIF) is a new framework for system identification in neuroscience. It integrates population receptive field estimation, neural encoding, connectivity analysis and hemodynamic response estimation in a single differentiable model that can be trained end-to-end via stochastic gradient descent. NIF models represent neural information processing systems as a network of coupled tensors, each encoding the representation of the sensory input contained in a brain region. The elements of these tensors can be interpreted as cortical columns whose activity encodes the presence of a specific feature in a spatiotemporal location. Each tensor is coupled to the measured data specific to a brain region via low-rank observation models that can be decomposed into the spatial, temporal and feature receptive fields of a localized neuronal population. Both these observation models and the convolutional weights defining the information processing within regions and effective connectivity between regions are learned end-to-end by predicting the neural signal during sensory stimulation. We trained a NIF model on the activity of early visual areas using a large-scale fMRI dataset. We show that we can recover plausible visual representations and population receptive fields that are consistent with empirical findings.

## 1 Introduction

Uncovering the nature of neural computations is a major goal in neuroscience [Churchland and Sejnowski, 1992]. It may be argued that a true understanding of the brain requires the development of *in silico* models that explain the behavior of biological neurons in terms of information processing. In cognitive terms, information processing can be understood as the use of internal representations of the environment to generate appropriate behaviour. A popular approach for uncovering the nature of these representations is to use a predefined set of stimulus features in order to predict measured neural responses to complex naturalistic stimuli in different brain regions [Naselaris et al., 2011, van Gerven, 2017, Yamins and DiCarlo, 2016]. Using this approach, the best results so far have been obtained using deep neural networks (DNNs) [Kriegeskorte, 2015, Güçlü and van Gerven, 2015b,a, Yamins and DiCarlo, 2016, Cichy et al., 2016, Horikawa and Kamitani, 2017, Cadena et al., 2019]. DNNs are machine learning algorithms that process the input through a sequence of layers of linear and nonlinear transformations. Each layer of a DNN encodes increasingly complex features of the original input. Strikingly, the hierarchy of features of DNNs has been shown to have a remarkable correspondence to the hierarchy of neural representations encoded in sensory brain regions [Güçlü and van Gerven, 2015b,a]. However, in these studies the DNNs were pretrained on arbitrary tasks such as object classification. Consequently, the resulting DNN features do not explicitly model the kind of computations that take place in individual neural populations and the predicted brain responses were obtained via a linear transformation of these pretrained features.

An alternative approach is to directly estimate parameterized neural models from measurements of neural activity. We refer to this approach as *neural system identification* [Stanley, 2005, Wu et al., 2006]. Neural system identification has been used to reveal mechanisms of neural information processing in biological systems [Joukes et al., 2014, Klindt et al., 2017, St-Yves and Naselaris, 2018, Antolík et al., 2016, McIntosh et al., 2016, Batty et al., 2017]. However, so far these ideas have mostly been applied within individual brain regions. More recent approaches learn to separate the location and features that voxels respond to, leveraging convolutional neural network training and visual cortex data [Klindt et al., 2017, St-Yves and Naselaris, 2018]. One aim of our method is to generalize these ideas under a common framework based on low-rank tensor observation models, with the potential of being applied beyond visual neuroscience. We expect that this framework will form the basis for analysis of upcoming large-scale data sets in neuroscience. Moreover, the behavior of complex networks of brain regions is usually studied using the methods of connectivity analysis and connectomics [Van Den Heuvel and Hulshoff Pol, 2010, Friston, 2011, Zuo et al., 2011, Sporns, 2013]. In particular, dynamic causal modeling (DCM) can be used to fit biologically detailed parameterized models of interacting brain regions [Friston et al., 2003]. Unfortunately, current applications of DCM do not allow to uncover the nature of information processing within and between brain regions since the activity of individual regions are described by a few scalar parameters that cannot be interpreted as a meaningful representation of the external stimuli.

This paper introduces a new framework, referred to as *neural information flow* (NIF), that combines the strengths of DNN-based neural encoding, neural system identification and connectivity analysis. Similar to prior brain encoding works, our framework extracts stimulus features from the layers of a DNN model. However, the layers of our DNN have a one-to-one correspondence to biological brain regions and all the model parameters are trained by fitting measured brain and behavioral responses. Each layer of the network encodes a family of spatially and temporally organized neural representations of the sensory input. In neurobiological terms, each unit of the layer can be interpreted as the activation of a cortical column that is responsive to a specific spatially localized feature such as an oriented bar in V1. These neural representations are obtained through convolutional layers that replicate the topologically organized connectivity between brain regions. In the case of functional magnetic resonance imaging (fMRI) analysis, the activity of the convolutional layers is coupled to the measured data through voxel-wise observation models that can be decomposed into a population receptive field, a hemodynamic response function and a channel (feature) receptive field. These observation models are trained jointly with the parameters of the network.

In the remainder of the paper we outline the basic principles of NIF in the context of a simplified model of visual information processing. However, the framework that we are introducing is general in the sense that the experimenter is free to choose the neural architecture of individual brain regions and how these regions map onto observed measurements, which can be either neural or behavioral in nature. The general philosophy of NIF is outlined in Figure 1. Given the generality of our framework, we expect that it will guide the development of a new family of computational models that allow us to uncover the principles of neural computations in biological systems. In the following, we outline the basic methodology of NIF modeling. Using a large fMRI dataset acquired under naturalistic stimulation we demonstrate that the model is capable of generating realistic brain measurements and that the computations captured in the model are biologically meaningful.

**Figure 1:**
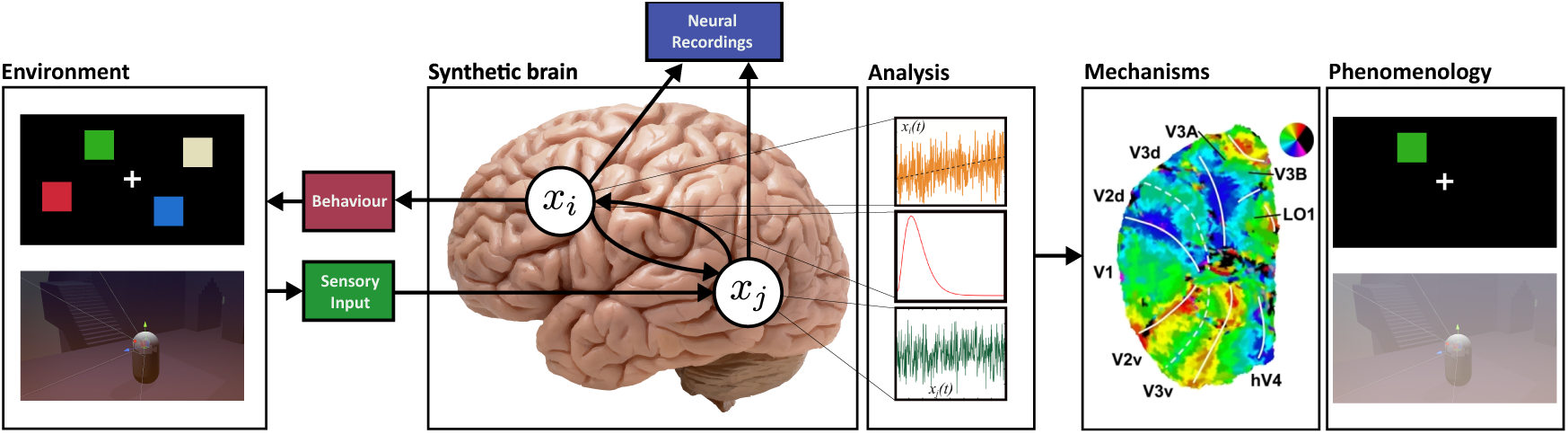
The philosophy underlying neural information flow. NIF models define synthetic brains that model information processing in real brains. They are specified in terms of mutually interacting neuronal populations (white discs) that receive sensory input (green) and give rise to measurements of neural activity (blue) and/or behavior (red). In practice, NIF models may consist of up to hundreds such interacting populations. They can be estimated by fitting them to neurobehavioral data acquired under these tasks. By analyzing NIF models, we can gain a mechanistic understanding of neural information processing in real brains and how neural information processing relates to phenomenology.

## 2 Neural information flow

The purpose of a NIF model is to capture the neural computations that take place within and between neuronal populations in response to sensory input. The core of a NIF model is a deep modular neural network architecture where individual neuronal populations are modeled using neural network modules that transform afferent input into efferent output. The connectivity between populations is captured by convolutional layers which model the topographically organized information exchange between neuronal populations. Finally, population activity is used to predict observed measurements through factorized observation models. Model parameters are estimated by fitting the neural signals measured during sensory stimulation. Specifically, the NIF model receives the same sensory input that is presented to the participant and predicts the measurements of all brain regions of interest. Model components are trained end-to-end using stochastic gradient descent (SGD); with the losses being the errors of these voxel-wise measurement predictions. In the following we describe the NIF components in more detail.

### 2.1 Modeling sensory input and neural representations

Sensory input is modeled using a four-dimensional tensor 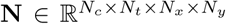 whose array dimensions represent input channels *c*, time *t* and spatial coordinates (*x, y*) respectively. For example, the input channels can be the RGB components of a visual stimulus or the photoreceptor responses of a retinal model. In our experiments, we model grayscale images using a single luminance channel (*N*_*c*_ = 1). We used temporal windows of 2.1s, resulting in 48 frames (*N*_*t*_ = 48). Analogously, the representations of the sensory input encoded in each brain region are modeled using four-dimensional tensors. The feature maps **N**[*c*, :, :, :] of these neural tensors encode neural processing of specific sensory stimulus features such as oriented edges or coherent motion. Consequently, a tensor element can be interpreted as the response of one cortical column. Under the same interpretation, cortical hyper-columns are represented by a sub-tensor **N**[:, :, *x, y*] storing the activations of all the columns that respond to the same spatial location.

### 2.2 Modeling directed connectivity and information flow

We model the directed connectivity between brain regions using spatiotemporal convolutions. The spatial weights model the topographically organized synaptic connections while the temporal component models synaptic delays.^1^ Using this setup, we can model how neural populations respond to sensory input as well as to each other.

Let **Ñ** denote the concatenation of afferent inputs **N**_1_, …, **N**_*N*_ along the feature dimension and let *\** denote the convolution operation. We define the activation of the *j*-th brain area as a function of its afferent input as follows:

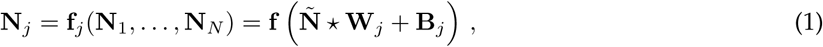

where **f** is the element-wise application of a sigmoid activation function followed by downsampling using an average pooling operation, **W**_*j*_ is a synaptic weight kernel and **B**_*j*_ is a bias term.

### 2.3 Modeling observable signals

NIF models are estimated by linking neural tensors to observation models that capture indirect measurements of brain activity. Observations are represented using tensors **Y** that store measurable responses. The observation model expresses the predicted measurements as a function of the activity of the latent tensors:

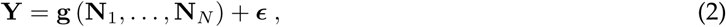

where ***ϵ*** is measurement noise. The exact form of **g** depends on the kinds of measurements that are being made. Neuroimaging methods such as fMRI, single- and multi-unit recordings, local field potentials, calcium imaging, EEG, MEGbut also motor responses and eye movements are observable responses to afferent input and can thus be used as a training signal. Note that the same brain regions can be observed using multiple observation models, conditioning them on multiple heterogeneous datasets at the same time. This provides a solution for multimodal data fusion in neuroscience [Uludağ and Roebroeck, 2014].

In this paper, we focus on modeling blood-oxygenation-level dependent (BOLD) responses obtained for individual voxels using fMRI. In this case, we can consider the voxel responses separately for each region, such that we have **Y**_*i*_ = **g**_*i*_ (**N**_*i*_) + ***ϵ*** for each region *i*. Let **Y**_*i*_ *∈* ℝ^*K*×*T*^ denote BOLD responses of *K* voxels acquired over *T* time points for the *i*-th region. Our observation model for the *k*-th voxel in that region is defined as

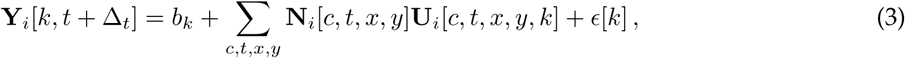

where *b*_*k*_ is a voxel-specific bias, *ϵ*[*k*] is normally distributed measurement noise and Δ_*t*_ is a temporal shift of the BOLD response that is used to take into account a default offset in the hemodynamic delay (4.9s in our experiments). Every brain region can be observed using a function of the form shown in Equation (3).

#### Factorized observation models

To simplify parameter estimation and facilitate model interpretability we use a factorized representation of **U**. That is,

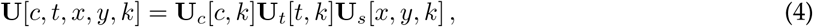

where **U**_*c*_[·, *k*] are the feature loadings that capture the sensitivity of a voxel to specific input features, **U**_*t*_[·, *k*] is the temporal profile of the observed BOLD response of the *k*-th voxel and **U**_*s*_[·, *·*, *k*] is the spatial receptive field of a voxel. Hence, the estimated voxel-specific observation models have a direct biophysical interpretation.

We further facilitate parameter estimation by using a spatial weighted low-rank decomposition of the spatial receptive field:

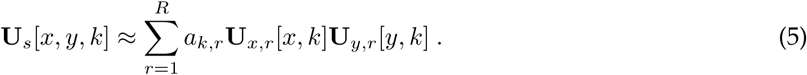

Here, *a*_*k,r*_ are rank amplitudes that are constrained to be positive using a softplus transformation. We used *R* = 4 in our experiments.^2^ To further stabilize the model and obtain localized population receptive fields, we apply a softmax nonlinearity to the columns of **U**_*t*_, **U**_*x*_ and **U**_*y*_. That is, the elements *u*_*i*_ of each column vector **u** of these matrices are given by

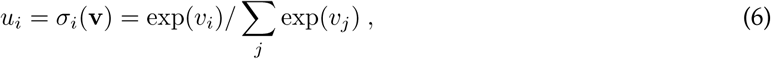

where the *v*_*i*_ are learnable parameters. This enforces positively weighted spatiotemporal receptive fields and reduces noise in the individual estimation of the voxel-specific weights.

### 2.4 Model estimation

Once the architecture of the NIF model is defined, synaptic weights and observation model parameters can be estimated by maximum likelihood using SGD. The individual loss terms for every brain region are summed to obtain a final loss that is minimized using SGD. Note that, since the model couples neuronal populations, region-specific estimates are constrained by one another and consequently make use of all observed data. The NIF example presented here was implemented in the `Chainer` framework for automatic differentiation [Tokui et al., 2015].

## 3 Experimental validation

To demonstrate the capabilities of the NIF framework, we estimated and tested a simple visual system model using a unique large-scale functional MRI dataset collected while one subject was exposed to almost 23 hours of complex naturalistic spatiotemporal stimuli. Specifically, we presented episodes from the BBC series *Doctor Who* [Davies et al., 2005].

### 3.1 Stimulus material

A single human participant (male, age 27.5) watched 30 episodes from seasons 2 to 4 of the 2005 relaunch of *Doctor Who*. This comprised the training set which was used for model estimation. Episodes were split into 12 min chunks (with each last one having varying length) and presented with a short break after every two runs. The participant additionally watched repeated presentations of the short movies *Pond Life* (five movies of 1 min, 26 repetitions) and *Space / Time* (two movies of 3 min, 22 repetitions), in random permutations after most episodes. They were taken from the series’ next iteration to avoid overlap with the training data. This comprised the test set which was used for model validation.

### 3.2 Data acquisition

We collected 3T whole-brain fMRI data. It was made sure that the training stimulus material was novel to the participant. Data were collected inside a Siemens 3T MAGNETOM Prisma system using a 32-channel head coil (Siemens, Erlangen, Germany). A T2*-weighted echo planar imaging pulse sequence was used for rapid data acquisition of whole-brain volumes (64 transversal slices with a voxel size of 2.4 × 2.4 × 2.4 mm^3^ collected using a TR of 700 ms). We used a multiband-multi-echo protocol with multiband acceleration factor of 8, TE of 39 ms and a flip angle of 75 degrees. The video episodes were presented on a rear-projection screen with the `Presentation` software package, cropped to 698 × 732 pixels squares so that they covered 20 degree of the vertical and horizontal visual field. The participant’s head position was stabilized within and across sessions by using a custom-made MRI-compatible headcast, along with further measures such as extensive scanner training. The participant had to fixate on a fixation cross in the center of the video. At the beginning of every break and after every test set video a black screen was shown for 14 s to record the fadeout of the BOLD signal after video presentation stopped. The black screen stimuli of these periods were omitted in the present analysis. In total this leaves us with 118.417 whole-brain volumes of single-presentation data, forming our training set (used for model estimation) and 1.032 volumes of resampled data, forming our test set (used for model evaluation). Data collection was approved by the local ethical review board (CMO regio Arnhem-Nijmegen, The Netherlands, CMO code 2014-288 with amendment NL45659.091.14) and was carried out in accordance with the approved guidelines. All specifics of the data set are described in a separate manuscript accompanying the data [Seeliger et al., 2019].

### 3.3 Data preprocessing

Minimal BOLD data preprocessing was performed using `FSL v5.0`. Volumes were first aligned within each 12 min run to their center volume (run-specific reference volume). Next, all run-specific reference volumes were aligned to the center volume of the first run (global reference volume). The run-specific transformations were applied to all volumes to align them with the global reference volume. The signal of every voxel used in the model was linearly detrended, then standardized (demeaning, unit variance) per run. Test set BOLD data was averaged over repetitions to increase signal to noise ratio, and as a final step the result was standardized again. A fixed delay of 7 TRs (4.9 s) was used to associate stimulus video segments with responses and allow the model to learn voxel-specific HRF delays within **U**_*t*_. With the video segments covering 3 TRs starting from the fixed delay, the BOLD signal corresponding to a stimulus is thus expected to occur within a time window of 4.9 s to 7.0 s after the onset of the segment. As there were small differences between frame rates in the train and test sets we converted the stimulus videos to a uniform frame rate of 23.86 Hz (16 frames per TR) for training the example model. To reduce model complexity we downsampled the videos to 112 × 112. As the model operates on three consecutive TRs, the training input size was 112 × 112 × 48. The stimuli were converted to grayscale prior to presenting them to the model. Otherwise stimuli were left just as they were presented in the experiment.

### 3.4 Model architecture

We implemented a purely feed-forward architecture for modeling parts of the visual system (LGN, V1, V2, V3, FFA and MT). The used architecture is illustrated in detail in Figure 2. FFA and MT have their own tensors originating from V3 to allow for a simplified model of the interactions between upstream and downstream areas. We intentionally used a simplified model to focus on demonstrating the capabilities of the NIF framework. To model LGN output, we used a linear layer consisting of a single 3 × 3 × 1 spatial convolutional kernel. The model was trained for 11 epochs with a batch size of 3, using the Adam optimizer [Kingma and Ba, 2014] with learning rate *α* = 5 × 10^−4^. Weights were initialized with Gaussian distributions scaled by the number of feature maps in every layer [He et al., 2015].

**Figure 2:**
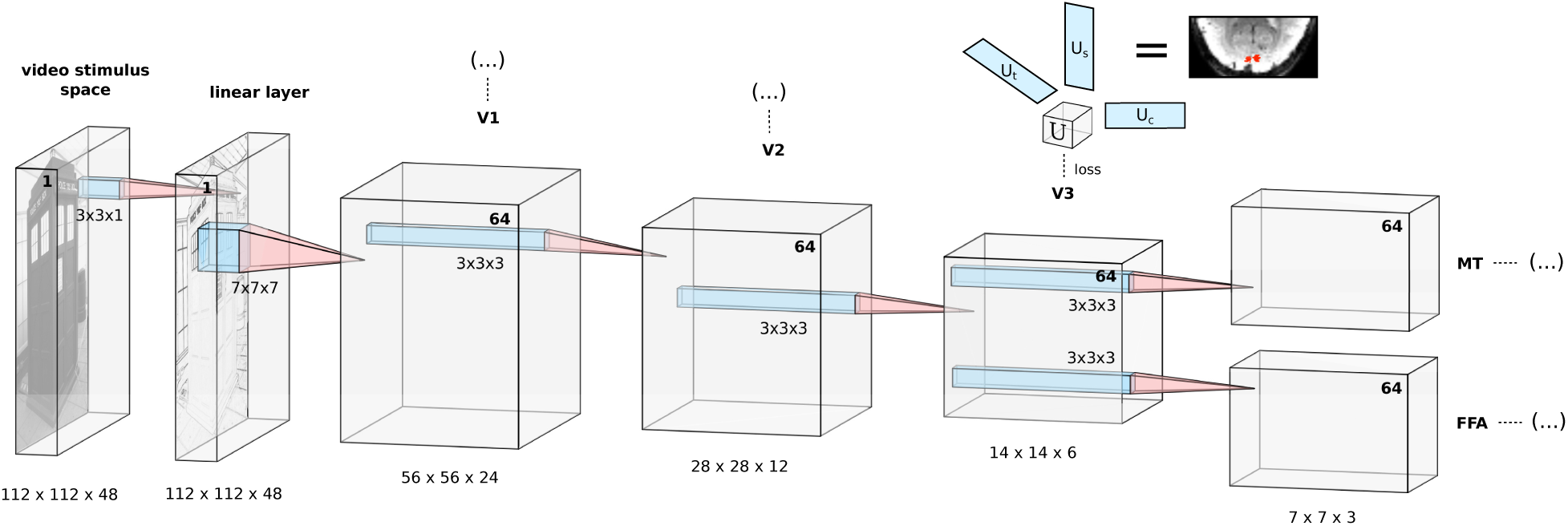
The employed NIF architecture providing a simplified feed-forward model of early visual areas. Underneath the tensors resulting from the 3D convolution operations we state the size of each input space (x × y × t) to the next layer. The number of feature maps in each input space is printed in boldface, with the stimulus (input) space consisting of a single channel. The input to the network are 3D stimulus video segments consisting of 3 × 16 frames (covering three TRs of 700 ms each), aligned with the hemodynamic response by applying a fixed delay of 7 TRs. The first convolutional layer is not attached to a region observation model, but is a single-channel linear spatial convolution layer. It serves as a learnable linear preprocessing step that accounts for retinal and LGN transformations. Convolutional kernel sizes are 7 × 7 × 7 in the second convolutional layer (leading to the V1 tensor), and 3 × 3 × 3 for all other layers. After every convolution operation we apply a sigmoid nonlinearity and spatial average pooling with 2 × 2 × 2 kernels. Before entering the **U**_t_ observation models the temporal dimension is average pooled so that each point t covers one TR. All weights in this model (colored blue) are learned by backpropagating the mean squared error losses from predicting the BOLD activity of the observed voxels. The voxel-specific observation models consisting of the spatiotemporal weight vectors **U**_s_ and **U**_t_ and the feature observation model **U**_c_ enable the end-to-end training of the model from observational data.

## 4 Results

In this paper we focus on the processing of visual information. In the following, we show that a NIF model uncovers meaningful characteristics of the visual system.

### 4.1 Accuracy of response predictions

After training the NIF model, we tested its accuracy on the test set. We observed that BOLD responses in a majority of voxels in each brain region could be significantly predicted by the model (*p <* 0.01, Bonferroni-corrected for the total number of gray matter voxels). This is illustrated in Figure 3, showing voxel-wise correlations between predicted and observed test data per region. The results show that the NIF model generates realistic brain activity in response to unseen input stimuli.

**Figure 3:**
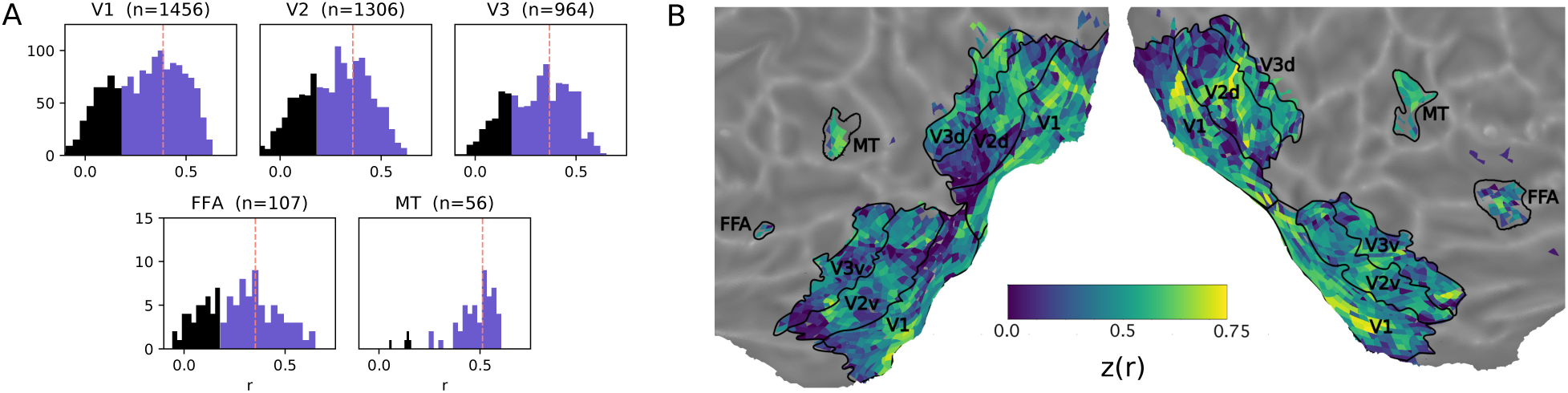
Voxel-wise correlations. A. Histograms of voxel-wise correlations between predicted and observed BOLD responses on the test set in different observed brain regions. The vertical line marks the median. The blue area shows the significantly predicted voxels. B. Cortical flatmap of the distribution of all correlations across the visual system. For the map we applied a Fisher z-transform to facilitate linear visual comparison of correlation magnitudes.

### 4.2 Visualization of learned representations

In this subsection we examin the features of the external stimulus that are encoded in our trained model of the visual system. We will begin with an analysis of the first layers, LGN and V1, whose features can be visualized by plotting the weights of the convolutional kernels. We will then show visualization of higher order regions using a more sophisticated preferred input analysis.

#### Linear feature analysis

For the first layers of the model, before the application of nonlinear transformations, neural network features can be inspected by visualizing the learned weights. A linear single-channel spatial layer was used to represent the transformation of the visual input at the retinal/LGN stage, before it enters visual cortex [Graham et al., 2006, Dan et al., 1996]. Figure 4A shows the estimated kernel as well as the resulting image transformation when applying this kernel to the input. As we can see, the linear kernel learns to perform edge extraction.^3^

**Figure 4:**
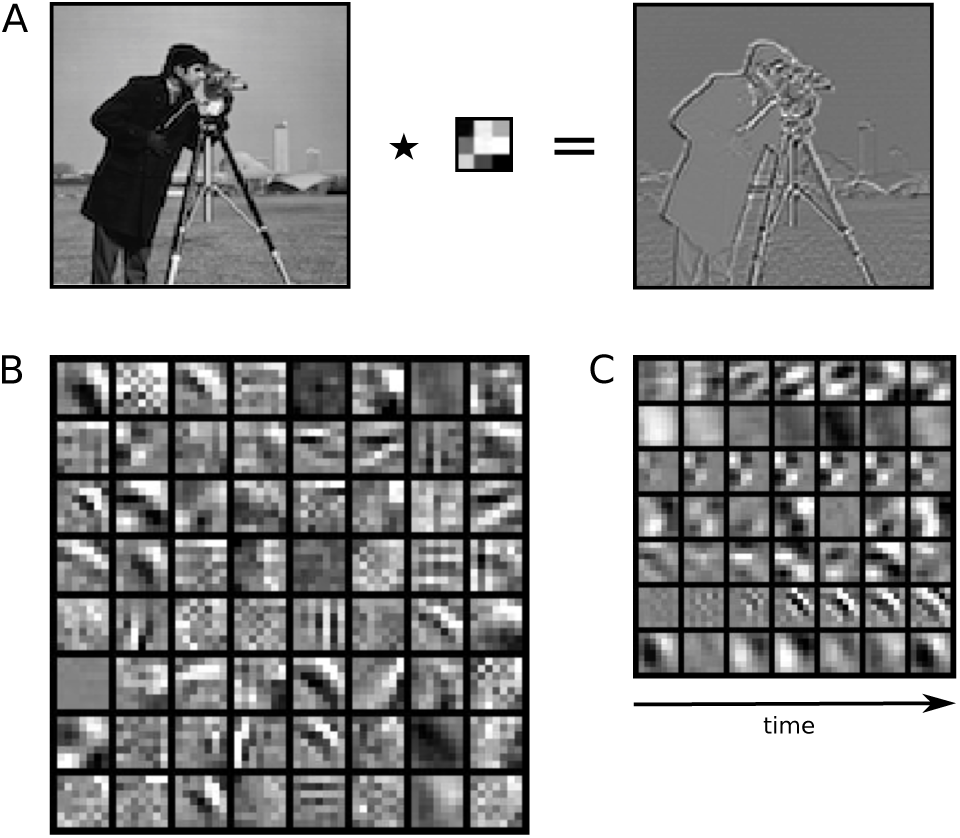
Stimulus features that were learned by the NIF model. A. Learned linear preprocessing showing that the estimated kernel extracts edges from the original input image. B. The 64 spatial features estimated from neural data for area V1 (frame three out of seven). C. Visualization of seven of these features across the temporal dimension. For visualization, feature weights were clipped at the extremes and all weights were globally rescaled between zero and one.

We can also visualize the feature detectors that determine the responses of V1. Figure 4B shows the 64 channels learned by the neural tensor connected to V1 voxels. Several well-known feature detection mechanisms of V1 arise, such as Gabor-like response profiles [Jones and Palmer, 1987]. As shown in Figure 4C, several of these feature detectors also show distinct dynamic temporal profiles, reflecting the processing of visual motion [Joukes et al., 2014].

#### Preferred input analysis

The linear feature analysis is insightful only for visualizing the features of LGN and V1. We can gain insight into the nature of the representations of higher order regions by visualizing which stimulus properties best drive simulated neural responses in a particular brain region. To this end, we estimated the gradient that leads to an increase in activity in individual target voxels, and used this gradient to modify the input such as to optimally drive the voxel response, starting from a three-dimensional white noise input. The technique is similar to [Bashivan et al., 2019]. The basic approach was originally proposed in [Erhan et al., 2009].

The analysis was performed only for those voxels for which the correlation between predicted and observed responses exceeded 0.4 on the test set. Let **y** = (*y*_1_, …, *y*_*K*_) such that **y** denotes the activity of all voxels that meet this predictability threshold in a specific ROI and *y*_*k*_ denotes the response of the *k*th target voxel. The objective is to optimize

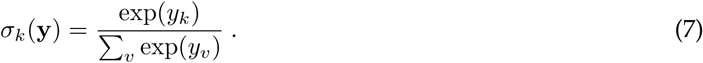

That is, we modify the input such as to maximize *y*_*k*_ while suppressing the responses of all other voxels in the same ROI *y*_*v*_ using a softmax nonlinearity. Let *I*_*t,x,y*_ denote the pixel intensity for the *t*th frame at spatial location (*x, y*). We further regularize the input using an *ℓ*_2_ loss on the components of *I* and a total variation loss *ℓ*_*TV*_ on neighboring timepoints and pixels in horizontal, vertical and diagonal directions. The objective is thus to minimize

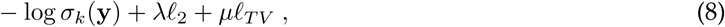

with *λ* = 10^−6^, *µ* = 10^−3^ for FFA and MT and *µ* = 5 · 10^−3^ in other ROIs. An SGD optimizer with an adaptive learning rate was used to optimize the stimuli. The iteration was stopped when no pixel changed more than 10^−3^ within 50 optimization steps. Optimization was done on a 112 × 112 × 16 video segment.

The results for different areas can be seen in Figure 5. V1 voxels show local Gabor patches at their respective spatial receptive fields. V2 and V3 show combinations of edge detectors consisting of multiple spatial frequencies. MT shows complex fields of varying small Gabor patches. FFA seems to prefer symmetry of simple geometric shapes.

**Figure 5:**
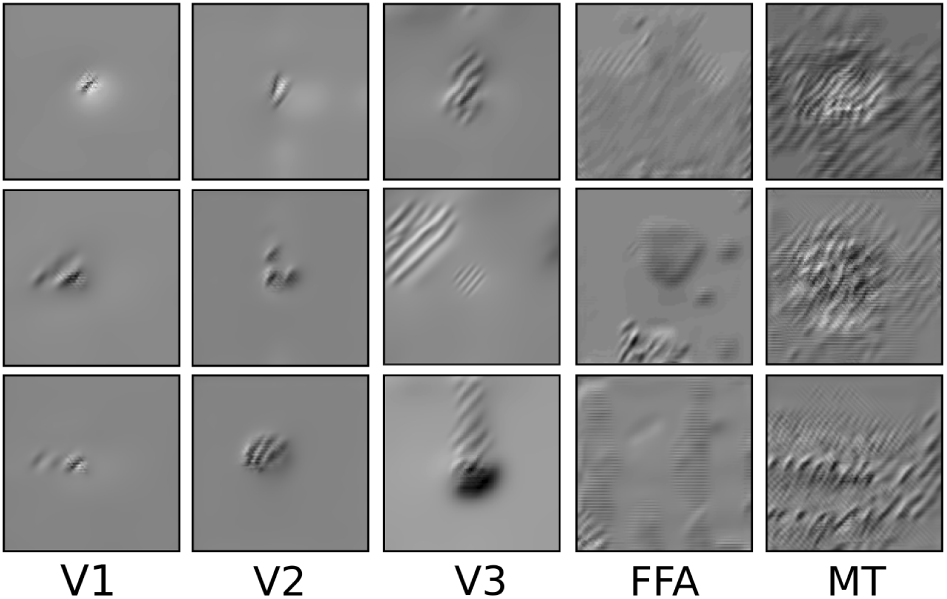
Examples of preferred inputs that maximize simulated voxel responses in different brain regions. Static frames. See supplementary material for the animated version.

### 4.3 Receptive field mapping

We examined whether the retinotopic organization of the visual cortex can be recovered from the spatial observation models [Wandell and Winawer, 2015]. Here, **U**_*s*_ directly represents spatial receptive field estimates for every voxel. Some of these voxel-specific receptive fields are shown in Figure 6A.

**Figure 6:**
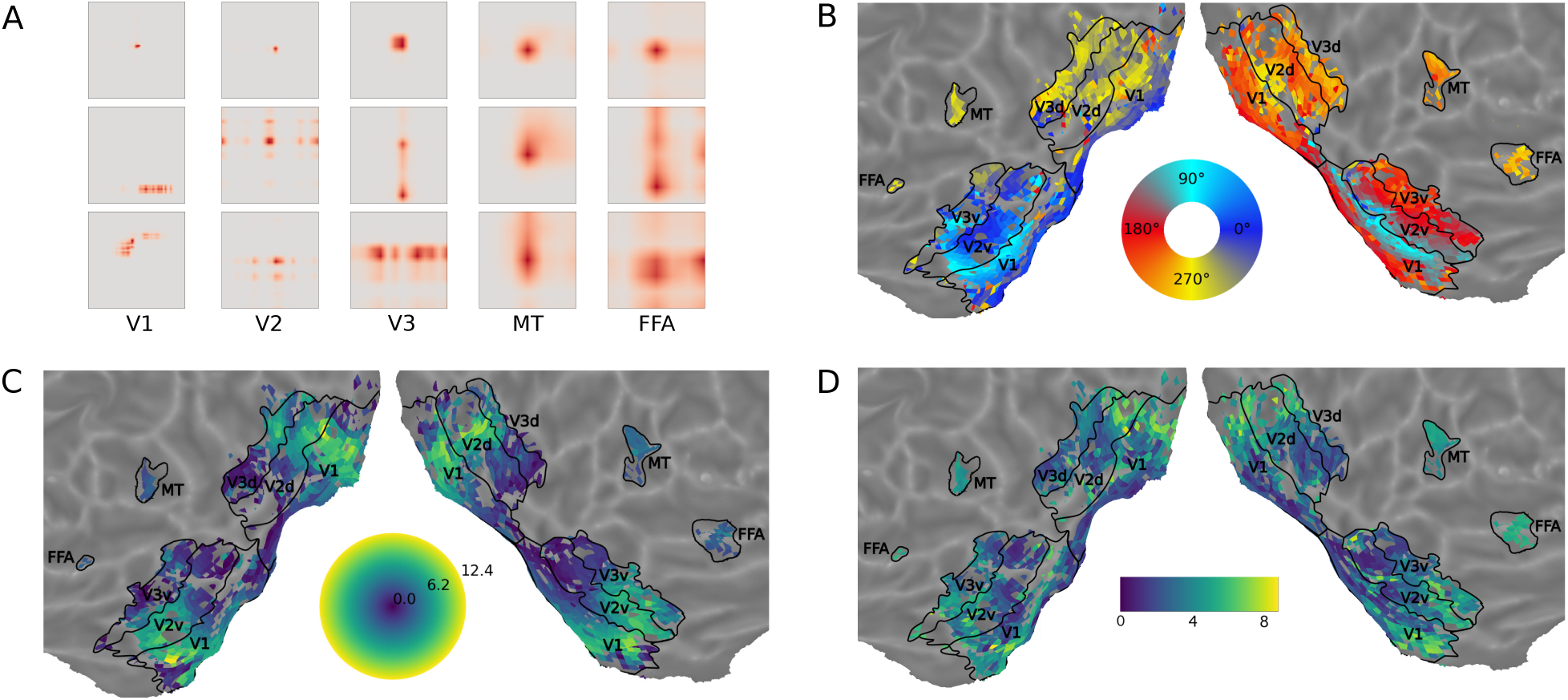
A. Various spatial receptive fields in video pixel space **U**_s_ learned for different ROIs within our framework. Most estimated spatial receptive fields are unipolar. B-D. Basic retinotopy that arose in the voxel-specific spatial observation matrix **U**_s_ within the NIF model. B. Polar angle. C. Eccentricity. D. Receptive field size.

To check that the NIF model has indeed captured sensible retinotopic properties, we determined the center of mass of the spatial receptive fields and transformed these centers to polar coordinates using the central fixation point as origin. Sizes of the receptive fields were estimated as the standard deviation across **U**_*s*_, using the centers of mass as mean. Voxels whose responses could not be significantly predicted were excluded from this analysis. Figure 6 shows polar angle (B), eccentricity (C) and receptive field size (D) for early visual system areas observed by our model. Maps were generated with `pycortex` [Gao et al., 2015]. Note that the boundaries between visual areas V1, V2 and V3 have been estimated with data from a classical wedge and ring retinotopy session (referred to as *wedge and ring localizers*). As can be seen, reversal boundaries align well with the traditionally estimated ROI boundaries. The larger eccentricity and increase in receptive field size (C) matches the expected fovea-periphery organization as well. Our results thus indicate that the NIF framework allows the estimation of accurate retinotopic maps from naturalistic videos.

### 4.4 Further properties of observational models

Recall that our model aims to predict the observed BOLD response from a spatiotemporal stimulus. We can obtain a rough estimate of the peak of the BOLD response by determining for each voxel the delay *t* that has the maximal weight **U**_*t*_[*t, k*] assigned. Figure 7 shows the distribution of these delays across cortex, providing an insight into spatial differences in the hemodynamic response function. Results show a consistent slowing of the HRF for downstream areas [Calhoun et al., 1998].

**Figure 7:**
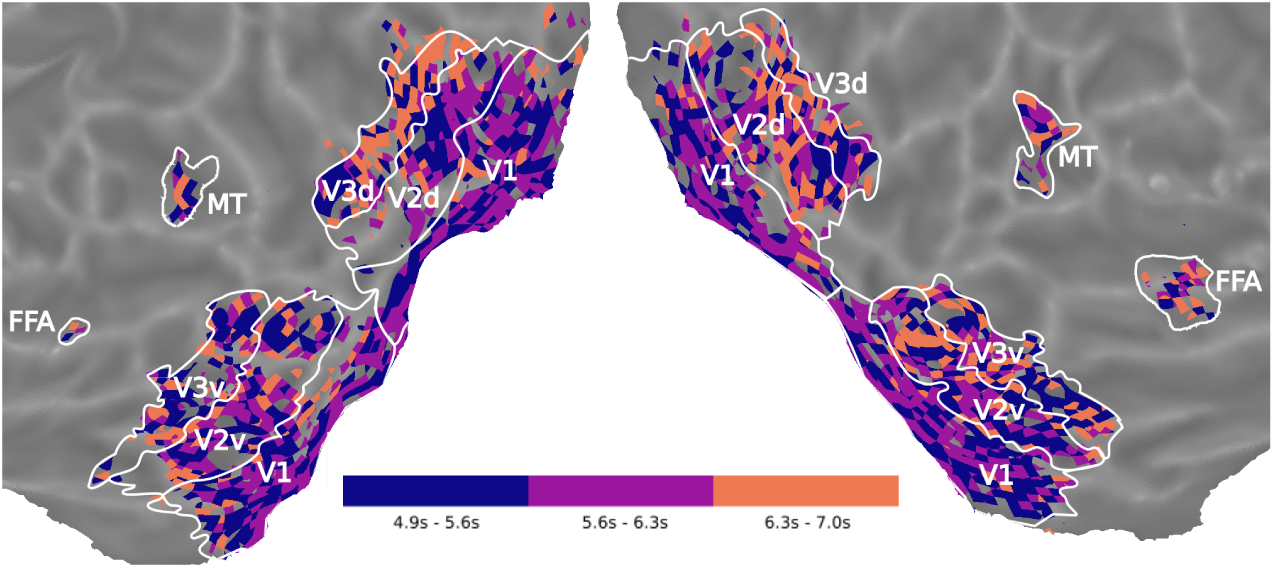
Differences in hemodynamic delay extracted from U_t_. For every voxel k we see the delay encoded in **U**_t_ [t, k] that has the maximal weight.

Furthermore, we can analyze **U**_*c*_. In Figure 8 we show the feature receptive field weight value for three different features in V1. We see different areas of early visual cortex showing inhibition or excitation for the selected features.

**Figure 8:**
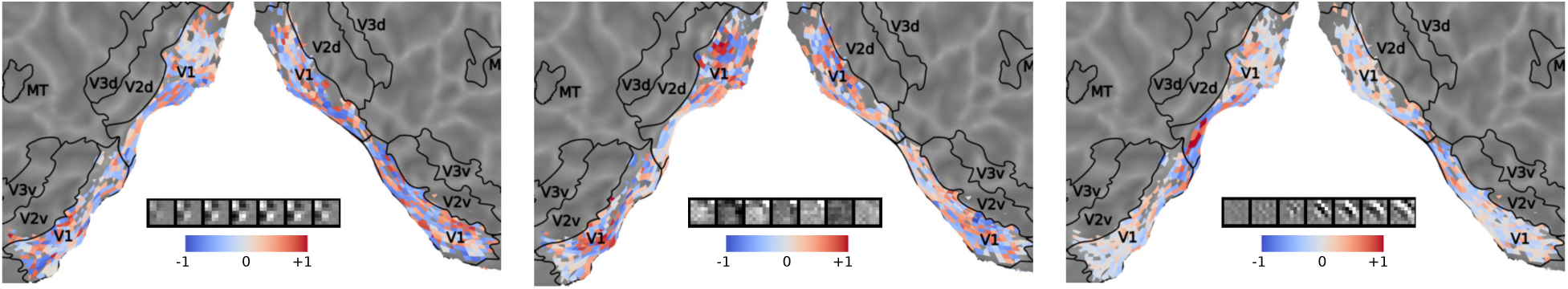
Projected U_c_ weight values for three different features in V1. Weight values were normalized between −1 and 1 by dividing them by the absolute maximum.

### 4.5 Processing of high-level semantic properties

So far, we have investigated characteristics of the NIF model that pertain to neural computations and representations and how these drive voxel responses. In this final analysis we investigate to what extent different neural populations are able to uncover high-level semantic content from the input stimulus. We focus on face detection since the processing of visual features pertaining to the discrimination of human faces is extremely well studied in the cognitive neuroscience literature [Haxby et al., 2000]. In particular, FFA is known to play a central role in the visual processing of human faces [Kanwisher et al., 1997]. Consequently, we expect that the representations learned by the FFA component of our model are related to human face processing.

We test this hypothesis using an *in silico* experiment closely resembling standard fMRI experimental procedures in cognitive neuroscience. We passed 90 2.1s long video segments, taken from the test set, through the trained NIF model. These videos were divided into two classes, one containing frontal views of human faces and the other not containing faces (45 videos per class). We analyzed the predicted BOLD responses of the models in the two experimental conditions using a mass univariate approach. For each voxel, we computed the t-statistic of the face minus no-face contrast and the associated p-values. We corrected for multiple comparisons using the false discovery rate (FDR) with alpha equal to 10^−4^. The left panel of Figure 9A shows the fraction of significant voxels in each brain region. The results show that FFA is the only region that is significantly activated by the contrast. The right panel shows that the voxels which are significantly activated also tend to be significantly predicted by the model. Figure 9B shows the significant (absolute) t-scores on the cortex.

**Figure 9:**
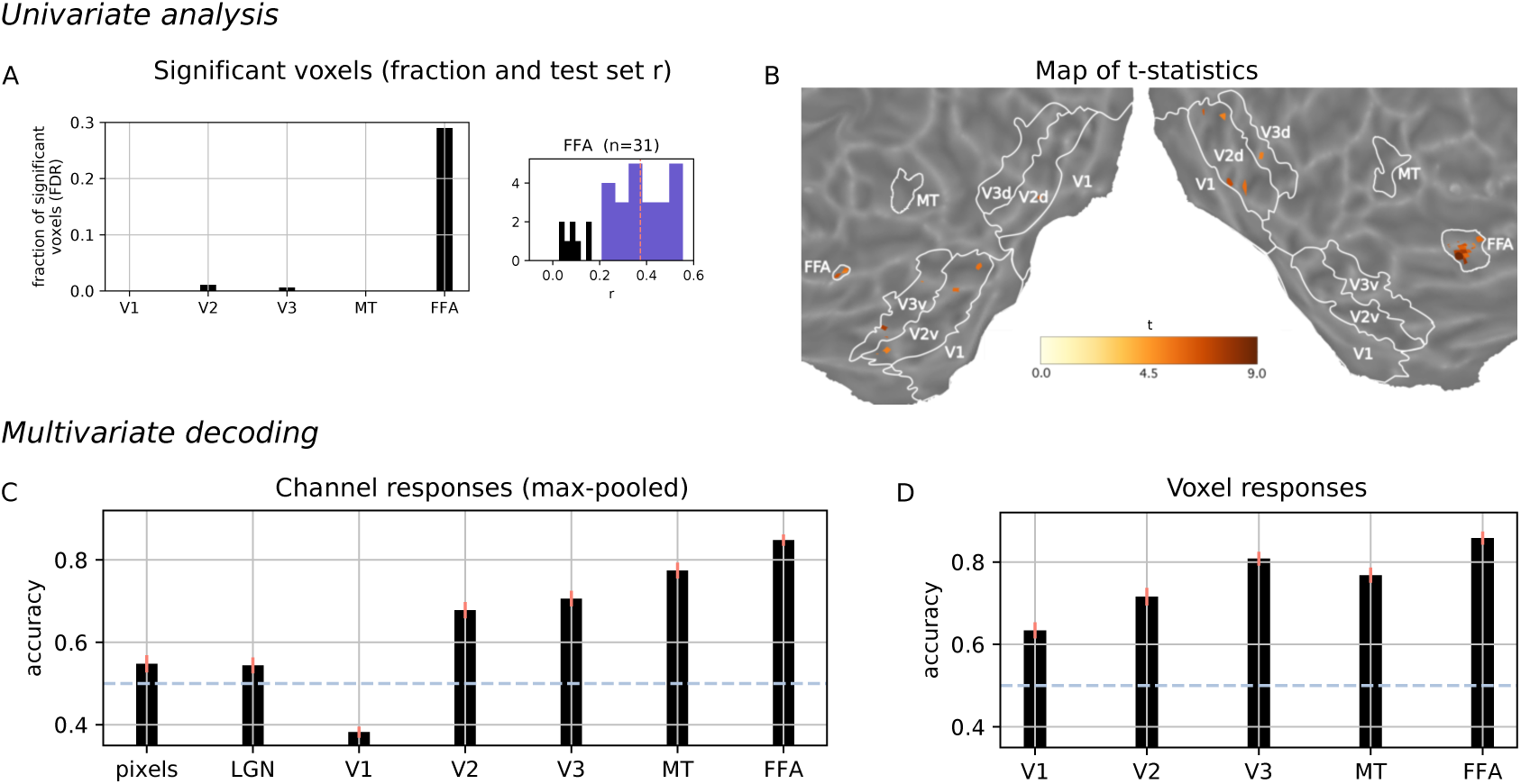
Results of an in silico experiment. The trained network was presented with video segments from the test set showing either faces or no faces. A., B. Univariate analysis. A. Significant voxels in each ROI. Correlations between predicted and observed voxel responses on the test set. B. Cortical map of the t-statistic for univariate analysis. C., D. Multivariate logistic regression. C. Decoding from ROI-wise tensor activations (channel responses max-pooled across the whole feature map) or raw input values (pixels, LGN). D. Decoding from predicted voxel responses. Overall, we see that FFA is the most discriminative area for the face recognition experiment.

We complemented these results with a multivariate decoding analysis [Haxby et al., 2014]. We trained a logistic regression model on the predicted voxel responses of each ROI in order to predict if the input contained faces. We also performed this logistic regression analysis directly on the channel responses of the model (max-pooled across the spatio-temporal feature map). In the analysis we also included direct predictions from the pixel values of the input images. We estimated the mean accuracy and its standard error by repeating the training 50 times with random splits into 35 training and 10 test examples respectively. As shown in Figure 9, the highest classification performance is achieved for FFA, both at the channel level and at the voxel level. This confirms our expectation that the model FFA has learned higher-order semantic properties that match its functional role in the brain. Furthermore, we see that multivariate data from increasingly downstream regions are more suitable to dissociate faces from non-faces. This indicates the prospect of studying *in silico* what behavioural goals higher-order sensory areas are optimized for. This also hints at the possibility of using neural information processing systems estimated from brain data to support the solution of pattern recognition tasks.

## 5 Discussion

This paper introduces neural information flow as a new approach for neural system identification. The approach relies on a neural architecture specified in terms of interacting brain regions, each performing nonlinear computations. By coupling each brain region with associated measurements of neural activity, we can estimate neural information processing systems end-to-end. We showed using fMRI data collected during prolonged naturalistic stimulation that we can successfully predict BOLD responses across different brain regions. Furthermore, meaningful receptive fields emerged after model estimation. The learned receptive fields are specific to each brain region but collectively explain all of the observed measurements. To the best of our knowledge, these results demonstrate for the first time that biologically interpretable information processing systems consisting of multiple interconnected brain regions can be directly estimated end-to-end from neural data.

As explained in the introduction, NIF generalizes current encoding models. For example, basic population receptive field models [Dumoulin and Wandell, 2008] and more advanced neural network models [van Gerven, 2017] are special cases of NIF that assume no interactions between brain regions and make specific choices for the nonlinear transformations that capture neuronal processing.

Our NIF approach is related to other recent approaches for neural system identification [Tripp, 2019, Klindt et al., 2017]. However, to the best of our knowledge, NIF models provide the first framework for estimating whole-brain neural information processing systems from data.

The present work also provides a new approach to connectivity analysis. The researcher can specify alternative NIF models and then use explained variance as a model selection criterion. This is similar in spirit to dynamic causal modeling. However, NIF models can identify changes in neural computation that are not detectable in models of causal interaction. For example, they can be used to investigate in detail the changes in neural information processing under different conditions.

NIF can be naturally extended in several directions. The employed convolutional layer to model neural computation can be replaced by neural networks that have a more complex architecture. For example, recurrent neural networks can be used to explicitly model the changes in neural dynamics that are now captured by 3D convolutions. Furthermore, lateral and feedback processing is easily included by adding additional links between brain regions and unrolling the backpropagation procedure over time. NIF models can also be extended to handle other data modalities. Alternative observation models can be formulated that allow us to infer neural computations from other measures of neural activity (e.g., single- and multiunit recordings, local field potentials, calcium imaging, EEG, MEG). Moreover, NIF models can be trained on multiple heterogeneous datasets at the same time, providing an elegant solution for multimodal data fusion. Our framework can also be easily applied to other sensory inputs. For example, auditory areas can be trained on auditory input (see e.g. [Güçlü et al., 2016]). If this is combined with visual input then we may be able to uncover new properties of multimodal integration [Simanova et al., 2014].

Furthermore, we are not restricted to using neural data as the sole source of training signal. We may instead (or also) condition these models on behavioral data, such as motor responses or eye movements. The resulting models should then show the same behavioral responses as the system under study. We can even teach NIF models to solve the task at hand directly using reinforcement learning [Sutton and Barto, 2017]. In this way, NIF models provide a starting point for creating brain-inspired AI systems that more closely model how real brains solve cognitive tasks.

Finally, we can use NIF models as *in silico* models to examine changes in neural computation. For example, we can examine how neural representations change during learning or as a consequence of virtual lesions in the network [Graziano and Aflalo, 2007]. This can provide insights into cognitive development and decline. We can also test what happens to neural computations when we directly drive individual brain regions with external input. This provides new approaches for understanding how brain stimulation modulates neural information processing, guiding the development of future neurotechnology [Roelfsema et al., 2018].

Summarizing, we view NIF as a starting point for building a new family of rich, general, biologically-inspired computational models that capture neural information processing in biological systems. As such, it provides a true blend of computational and experimental neuroscience [Churchland and Sejnowski, 2016]. NIF models are also scalable since they make use of efficient stochastic gradient methods, as developed by the artificial intelligence community. This gives us a principled approach to make sense of the high-resolution datasets produced by continuing advances in neurotechnology [Stevenson and Kording, 2011]. We expect that NIF models will deliver exciting new insights into the principles and mechanisms that determine neural information processing in biological systems.

## Acknowledgements

This research was supported by VIDI grant number 639.072.513 of The Netherlands Organization for Scientific Research (NWO).

## Footnotes

1 To enforce causality of the neural responses, the temporal filters should be causal, meaning that the only non-zero weights correspond to past time points. However, this assumption can be dropped when the time scale of our observations is much slower than that of the underlying temporal dynamics.

2 The rank limits the complexity of the spatial observation model. Rank one models can estimate unimodal receptive fields. However, a small number of voxels have nonclassical receptive fields that respond to multiple parts of the input space, for which more degrees of freedom are needed.

3 When using two kernels, the second kernel effectively learns to extract image luminance. This is likely to be a reflection of the independence of luminance and contrast information in natural images and in LGN responses [Mante et al., 2005].

